# Loss of phosphatidylserine flippase β-subunit *Tmem30a* in podocytes leads to albuminuria and glomerulosclerosis

**DOI:** 10.1101/2020.12.04.412635

**Authors:** Wenjing Liu, Lei Peng, Wanli Tian, Yi Li, Ping Zhang, Kuanxiang Sun, Yeming Yang, Xiao Li, Guisen Li, Xianjun Zhu

**Affiliations:** Sichuan Provincial Key Laboratory for Human Disease Gene Study, Sichuan Provincial People’s Hospital, University of Electronic Science and Technology of China, Chengdu, Sichuan, China; Department of Nephrology, Sichuan Academy of Medical Sciences & Sichuan Provincial People’s Hospital, Chengdu, Sichuan Clinical Research Center for Kidney Diseases, Sichuan, China; Department of Laboratory Medicine, Sichuan Academy of Medical Sciences & Sichuan Provincial People’s Hospital, Chengdu, Sichuan, China

## Abstract

Phosphatidylserine (PS) is asymmetrically concentrated in the cytoplasmic leaflet of eukaryotic cell plasma membranes. This asymmetry is regulated by a group of P4 ATPases (named PS flippases) and its β-subunit TMEM30A. The disruption of PS flippase leads to severe human diseases. *Tmem30a* is essential in the mouse retina, cerebellum and liver. However, the role of *Tmem30a* in the kidney, where it is highly expressed, remains unclear. Podocytes in the glomerulus form a branched interdigitating filtration barrier that can prevent the traversing of large cellular elements and macromolecules from the blood into the urinary space. Damage to podocytes can disrupt the filtration barrier and lead to proteinuria and podocytopathy, including focal segmental glomerulosclerosis, minimal change disease, membranous nephropathy, and diabetic nephropathy. To investigate the role of *Tmem30a* in the kidney, we generated a podocyte-specific *Tmem30a* knockout (cKO) mouse model using the NPHS2-Cre line. *Tmem30a* KO mice displayed albuminuria, podocyte degeneration, mesangial cell proliferation with prominent extracellular matrix accumulation and eventual progression to focal segmental glomerulosclerosis (FSGS). Reduced *TMEM30A* expression was observed in patients with minimal change disease and membranous nephropathy, highlighting the clinical importance of TMEM30A in podocytopathy. Our data demonstrate a critical role of *Tmem30a* in maintaining podocyte survival and glomerular filtration barrier integrity. Understanding the dynamic regulation of the PS distribution in the glomerulus provides a unique perspective to pinpoint the mechanism of podocyte damage and potential therapeutic targets.

## Introduction

Phosphatidylserine (PS) is asymmetrically and dynamically distributed across the lipid bilayer in eukaryotic cell membranes [1]. Such dynamic distribution is preserved by flippases, one of the most important P4-ATPases, which possess flippase activity that catalyses lipid transportation from the outer to the inner leaflet to generate and maintain phospholipid asymmetry [2]. The PS asymmetry maintained by P4-ATPases is essential to various cellular physiological and biochemical processes, including vascular trafficking, cell polarity and migration, cell apoptosis and cell signalling events [2-6].

As the β-subunit of P4-ATPases (except ATP9A and ATP9B), TMEM30 family proteins play essential roles in the proper folding and subcellular localization of P4-ATPases [7, 8]. The TMEM30 (also called CDC50) family includes TMEM30A, TMEM30B and TMEM30C, of which TMEM30A interacts with 11 of the 14 P4-ATPases [9-13]. Our previous studies have demonstrated that TMEM30A deficiency causes a series of disorders: retarded retinal angiogenesis, Purkinje cell, retinal bipolar cell and photoreceptor cell degeneration, impaired foetal liver erythropoiesis, intrahepatic cholestasis and chronic myeloid leukaemia [5, 14-19].

The glomerular filtration barrier includes three layers: fenestrated endothelial cells, the glomerular basement membrane (GBM) and glomerular epithelial cells (podocytes). Podocytes consist of a cell body that gives rise to major processes and minor foot processes (FPs). The FPs of neighbouring podocytes form a branched interdigitating network, and the space between adjacent FPs is covered by a multiprotein complex called the slit diaphragm (SD), the final barrier [20]. The glomerular filtration barrier prevents the traversing of large cellular elements and macromolecules from the blood into the urinary space, and defects in the selective barrier result in albuminuria and nephrotic syndrome. Damage to podocytes can disrupt the filtration barrier, which is a key step of proteinuria and podocytopathy (including focal segmental glomerulosclerosis (FSGS), minimal change disease (MCD), membranous nephropathy (MN), and diabetic nephropathy (DN)), as well as other types of kidney diseases (such as immunoglobin A nephropathy (IgAN) and lupus nephritis). FSGS is one of the most widely used disease models to study podocytopathy and proteinuria [21].

Given that *Tmem30a* is essential for tissues with high TMEM30A expression, such as retina, cerebellar and hepatic tissue, and that *Tmem30a* is highly expressed in the kidney, we set out to elucidate the role of *Tmem30a* in the kidney by generating a podocyte-specific *Tmem30a* knockout (KO) model. *Tmem30a* KO mice displayed albuminuria, podocyte injury and loss, mesangial cell proliferation with prominent extracellular matrix (ECM) accumulation and eventual progression to FSGS. Furthermore, we observed markedly diminished *TMEM30A* expression in patients with MCD and MN, highlighting the clinical importance of TMEM30A in podocytopathy. Taken together, our findings demonstrate that *Tmem30a* plays a critical role in maintaining podocyte survival and glomerular filtration barrier integrity.

## Materials and Methods

### Mouse model

All animal protocols were approved by the Ethics Committee of Sichuan Provincial People’s Hospital. All animal experiments were performed according to the approved protocols and related guidelines. Mice were raised under a 12-h light/12-h dark cycle.

A conditional knockout (cKO) allele carrying a floxed *Tmem30a* allele (*Tmem30a^loxp/loxp^*) has previously been described [15-17]. To generate mice with *Tmem30a* deletion specifically in podocytes, *Tmem30a ^loxP/loxP^* mice were crossed with transgenic mice expressing Cre recombinase under the control of the podocyte-specific podocin (NPHS2) promoter (podocin-Cre, B6.Cg-Tg(NPHS2-cre)295Lbh/J, stock no.: 008205) [22] to yield progeny with the genotype of *Tmem30a^loxP/+^*; N PHS2-Cre. Cre-positive heterozygous offspring were crossed with *Tmem30a^loxP/loxP^* mice to obtain *Tmem30a^loxP/loxP^*; NPHS2-Cre cKO mice. A tdTomato reporter was introduced to monitor the efficiency of Cre-mediated deletion of the floxed exon (strain name: B6. Cg-Gt(ROSA)26Sortm14(CAG-tdTomato)Hze/J; Jackson Laboratory, stock no. 007914; http://jaxmice.jax.org/strain/007914.html). The reporter contains a loxP-flanked STOP cassette that prevents transcription of the downstream CAG promoter-driven red fluorescent protein variant tdTomato. In the presence of Cre recombinase, the STOP cassette is removed from the Cre-expressing tissue(s) in reporter mice, and tdTomato will be expressed.

### Genotyping by PCR

Genomic DNA samples obtained from mouse tails were genotyped using PCR to screen for the floxed *Tmem30a* alleles using primers for *Tmem30a*-loxP2-F, ATT CCC CTC AAG ATA GCT AC, and *Tmem30a*-loxP2-R, AAT GAT CAA CTG TAATTC CCC. Podocin-Cre was genotyped using generic Cre primers: Cre-F, TGC CAC GAC CAA GTG ACA GCA ATG, and Cre-R, ACC AGA GAC GCA AAT CCATCG CTC. TdTomato mice were genotyped using the following primers provided by the JAX mouse service: oIMR9020, AAG GGA GCT GCA GTG GAG TA; oIMR9021, CCG AAA TCT GTG GGA AGT C; oIMR9103, GGC ATT AAA GCA GCG TAT CC; and oIMR9105, CTG TTC CTG TAC GGC ATG G. The first cycle consisted of 95°C for 2 minutes, followed by 33 cycles of 94°C for 15 seconds, 58°C for 20 seconds and 72°C for 30 seconds.

### Urine analysis

Twenty-four-hour urine samples were collected using metabolic cages. Collected urine samples were centrifuged at 500 g for 5 min, and the supernatant was used for the quantitation of albumin and creatinine. Quantitation of urinary albumin and creatinine was carried out using mouse albumin-specific ELISA kits (Roche) and creatinine determination kits (Enzymatic Method) (Roche), respectively, following the manufacturer’s instructions.

### Renal pathology

Mice were anaesthetized with a combination of ketamine (16 mg/kg body weight) and xylazine (80 mg/kg body weight) and perfused transcardially with ice-cold PBS, followed by 4% paraformaldehyde in 100 mM PBS (pH 7.4). The kidneys were harvested, fixed in 4% paraformaldehyde, dehydrated and embedded in paraffin or optimal cutting temperature (OCT) solution for cryosectioning by standard procedures. Sections (2 µm) to be used for light microscopy analysis were subjected to periodic acid-Schiff (PAS) staining and visualized with a light microscope (Nikon Eclipse Ti-sr).

### Patient recruitment and ethics statement

The patient study was approved by the institutional review board of the Sichuan Provincial People’s Hospital in Chengdu, China. All experiments were carried out in accordance with the approved study protocol. All subjects enrolled signed written informed consent forms. Kidney tissues from IgAN, DN, MCD and MN patients were collected during renal biopsy in the Nephrology Department of Sichuan Provincial People’s Hospital, and adjacent normal renal tissues were collected from patients with renal tumours during nephrectomy in the Department of Urology at the same hospital. All human kidney tissues underwent routine renal pathological examination to confirm the diagnosis. These tissues were processed by standard procedures for cryosectioning and immunofluorescent staining, as described below.

### Immunohistochemistry and immunofluorescence

Paraffin-embedded murine kidney slides (2 µm) were deparaffinized following a standard protocol. After washing and blocking, the tissues were incubated with primary antibodies against Wilms tumour-1 (WT1) (1:100, Servicebio, GB11382) and synaptopodin (1:100, ZEN BIO, 508484). The slides were then incubated with HRP-labelled donkey anti-rabbit secondary antibodies. Nuclei were visualized using DAPI counterstaining. Glomerular WT1 was determined by counting positively immunostained nuclei in 30 glomerular profiles in each kidney section. Images were taken using a Zeiss Axioplan-2 imaging microscope with the digital image-processing program AxioVision 4.3.

Frozen mouse tissues were sectioned at 5 µm (CryoStar NX50 OP, Thermo Scientific, Germany). After blocking and permeabilization with 10% normal goat serum and 0.2% Triton X-100 in PBS at room temperature for 1 h, the cryosections were labelled with the following primary antibodies overnight at 4°C: TMEM30A (1:50; mouse monoclonal antibody Cdc50-7F4, gift from Dr Robert Molday, University of British Columbia, Canada) and nephrin (1:100, Abcam, Cambridge, MA, USA). The sections were rinsed in PBS three times and incubated with Alexa Fluor 488- or Alexa Fluor 594-labelled goat anti-mouse (Bio-Rad Laboratories, catalogue # STAR132P, RRID: AB_2124272) or anti-rabbit IgG secondary antibodies (diluted 1:500, Bio-Rad Laboratories, 5213-2504 RRID: AB_619 907), and then stained with DAPI at room temperature for 1 h. Images were captured on a laser scanning confocal microscope (LSM800, Zeiss, Thornwood, NY, USA).

Frozen human tissues were sectioned using a cryomicrotome (MEV, SLEE, Germany) at 4 µm. To observe the expression of TMEM30A, cryosections were stained with rabbit anti-human TMEM30A (1:100, Bioss, Beijing, China) overnight at 4°C followed by FITC-conjugated goat anti-rabbit IgG (1:100, Gene Tech Company Limited, Shanghai, China) at 37°C for 30 min. Images were captured using an Olympus BX51 microscope (Tokyo, Japan). All exposure settings were kept the same. The fluorescence intensity was measured by manually outlining the perimeter of every glomerulus and semiquantifying the luminosity of the outlined regions using image analysis software (ImageJ, version 1.52p, National Institutes of Health, USA). A background correction was made for each glomerulus by subtracting the average intensity in non-stained regions (outlined manually) in the glomerulus.

### Transmission electron microscopy (TEM)

TEM was performed on kidney cortical tissue (HITACHI, HT7700). Kidneys obtained from WT and KO mice were cut into small pieces just after harvest and fixed in fixative solution (2.5% glutaraldehyde, 1.25% paraformaldehyde, and 0.003% picric acid in 0.1 M sodium cacodylate buffer [pH 7.4]) for 2 h at room temperature. The fixed kidney was washed with 0.1 M PBS, postfixed with 1% osmium tetroxide (OsO4) in 0.1 M PBS (pH 7.4), and washed in 0.1 M phosphate buffer (pH 7.4) three times. The fixed tissue was embedded in Epon 812 after dehydration via an ascending series of ethanol and acetone and incubated at 60°C for 48 h. Ultrathin sections (60 nm) were cut and stained with uranyl acetate and lead citrate.

#### Isolation of Glomeruli

The glomeruli were dissected using standard sieving technique [23]. Briefly, kidney were mashed with syringe plunger and then pushed through 425 µm (top), 250 µm, 175 µm, 125µm, 100 µm and 70µm (bottom) sieve with ice cold mammalian Ringer’s solution (Shyuanye Biotechnology, Shanghai, China L15O10G100158) with 1% BSA (Solarbio, Beijing, China. A8010). Remove the top sieve and proceed to do the same on the next. Collect the glomerular retained by the 100 µm and 70µm sieve into centrifuge tube with ice cold mammalian Ringer’s solution with 1% BSA. Centrifuge the tube at 1,000g for 10 min at 4°C, remove the supernatant and then freeze the glomeruli in liquid N2 before storing at ™80°C for further protein and RNA extraction.

### Western blotting

Isolated glomerular proteins were extracted in RIPA lysis buffer (50 mM Tris-HCl, 150 mM NaCl, 1% Triton X-100, 0.5% sodium deoxycholate, and 0.1% SDS, pH 7.4) supplemented with complete protease inhibitor cocktail (Roche). The protein concentration was determined with the Bicinchoninic Acid (BCA) Protein Assay (Thermo Fisher). SDS-PAGE and Western blot analysis were performed with equal amounts of protein (15 µg), which were then transferred to PVDF membranes (GE Healthcare, Chicago, IL, USA). After blocking with 8% non-fat dry milk in TBST for 2 h at room temperature, the blots were probed with primary antibodies against CHOP (1:1000, Cell Signaling Technology, Danvers, MA, USA), BiP (1:1000, Cell Signaling Technology Danvers, MA, USA) and PDI (1:2000, Cell Signaling Technology, Danvers, MA, USA) in blocking solution overnight at 4°C, followed by incubation with anti-mouse or anti-rabbit HRP-conjugated secondary antibodies (1:5000, Cell Signaling Technology, Danvers, MA, USA). The samples were normalized with GAPDH (1:5000, Proteintech, Wuhan, China) primary antibody, and the relative intensity of the blots was quantified using ImageJ software.

### Statistical analysis

Data are expressed as the mean ± standard error of the mean (SEM). Statistical evaluation was performed using Student’s *t* test. P values of <0.05 were considered to be statistically significant.

## Results

### Generation of podocyte-specific *Tmem30a* KO mice

Previous studies have demonstrated the essential role of *Tmem30a* in several vital tissues. In the retina, *Tmem30a* is important for the survival of retinal photoreceptor and rod bipolar cells [16, 17]. In the cerebellum, *Tmem30a* loss results in early-onset ataxia and cerebellar atrophy [15]. In the liver, *Tmem30a* deficiency impairs mouse foetal liver erythropoiesis and causes intrahepatic cholestasis by affecting the normal expression and localization of bile salt transporters and causes intrahepatic cholestasis [5, 18]. In the haematopoietic system, *Tmem30a* is critical for the survival of haematopoietic cells and leukocytes [19]. *Tmem30a* is expressed in the retina, brain, cerebellum, liver, heart, kidney, spine, and testis [8, 17, 23], but its role in the kidney remains elusive. To define the role of *Tmem30a* in the kidney, we first assessed the expression of *Tmem30a* in the kidney by immunostaining with a proven TMEM30A antibody [17]. Kidney cryosections were immunostained with specific antibodies against *Tmem30a* (Fig. 1a). *Tmem30a* is highly expressed in the glomeruli, which implies a vital role of *Tmem30a* in glomerular filtration. To investigate this role of *Tmem30a*, we generated podocyte-specific *Tmem30a* KO *Tmem30a*^loxP/loxP^; Nphs2-Cre (hereafter named *Tmem30a* KO) mice by crossing *Tmem30a*^loxP/loxP^ with podocin-cre Nphs2-Cre mice (Fig. 1b). *Tmem30a* expression was reduced by ~55% in the glomerulus of *Tmem30a* KO mice compared with that in control mice (Fig. 1c). Given the presence of Cre only in the podocytes, the deletion efficiency was fairly good. ROSA26-tdTomato was used to verify the specific expression of podocin-cre in podocytes. We crossed *Tmem30a*^loxP/+^; Nphs2-Cre; Rosa-tdTomato mice with *Tmem30a*^loxP/loxP^ mice to generate littermate *Tmem30a*^+/+^; NPHS2-Cre; Rosa-tdTomato and *Tmem30a*^loxp/loxp^; NPHS2-Cre; Rosa-tdTomato mice to evaluate the KO specificity of *Tmem30a* in podocytes (Fig. 1b-d). In summary, these data demonstrate the successful elimination of *Tmem30a* in *Tmem30a*^loxp/loxp^; NPHS2-Cre mice.

**Fig. 1.**
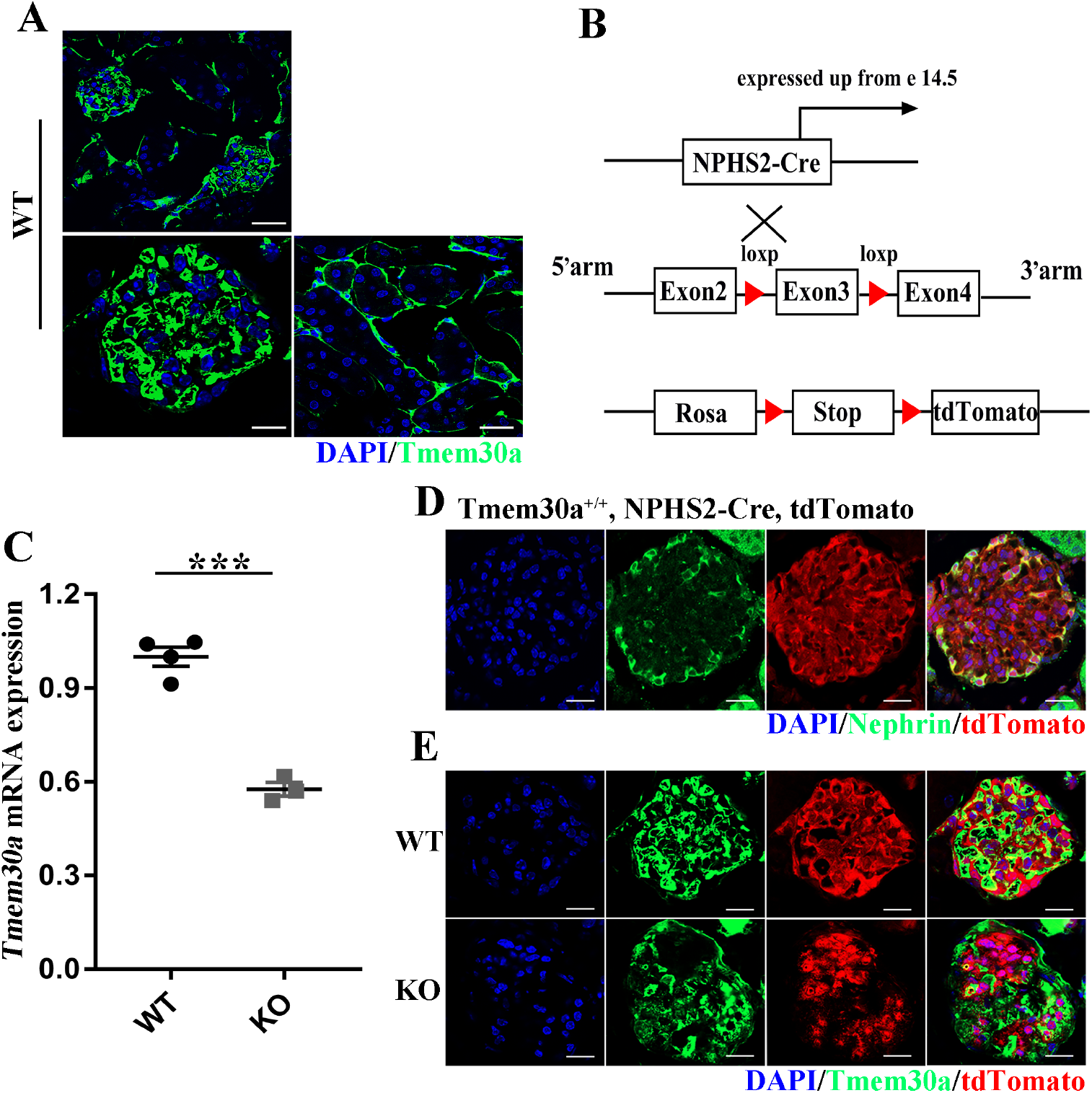
Generation of podocyte-specific *Tmem30a* cKO mice. (A) Cryosections of the kidney from 5-month-old WT mice were immunostained with *TMEM30A* antibody (green). The upper panel provide a lower magnification *TMEM30A* staining image of the glomerular cortex and the lower panel shows high resolution immunostaining of the glomeruli and renal tubules for the *TMEM30A* protein, respectively. *Tmem30a* is highly expressed in the glomeruli (scale bar: the upper panel: 25µm; the lower panel: 10µm). (B) Schematic showing the targeting strategy for generating podocyte-specific *Tmem30a*-KO mice. Rosa-tdTomato reporter mice were used to monitor Cre expression. (C) Q-PCR showed the relative mRNA expression of *Tmem30a* in the glomeruli of KO mice compared with those of WT mice. Sample size, n=4. (D) The ROSA-tdTomato reporter was introduced to monitor the expression of Cre recombinase (red). Podocytes were labelled with the podocyte-specific marker nephrin (green). TdTomato-expressing cells were colocalized with nephrin-labelled podocytes, indicating the specific expression of NPHS2-Cre. (E) Localization of *TMEM30A* and Rosa-tdTomato in WT and KO mice by immunofluorescence, suggesting that *TMEM30A* was knocked out in podocytes (scale bar, 25 µm).

### Podocyte-specific deletion of *Tmem30a* results in albuminuria

*Tmem30a* KO mice were born at the ratio that is consistent with classic Mendelian segregation. No obvious morphological abnormalities were observed in *Tmem30a* KO mice upon gross examination. Although they appeared to be normal in terms of body size, the albuminuria level in *Tmem30a* KO mice increased significantly compared with control mice from 5 months after birth (Fig. 2). By the ninth months after birth, the albuminuria level continued to rise, indicating sustained impairment of the glomeruli and selective barrier (Fig. 2).

**Fig. 2.**
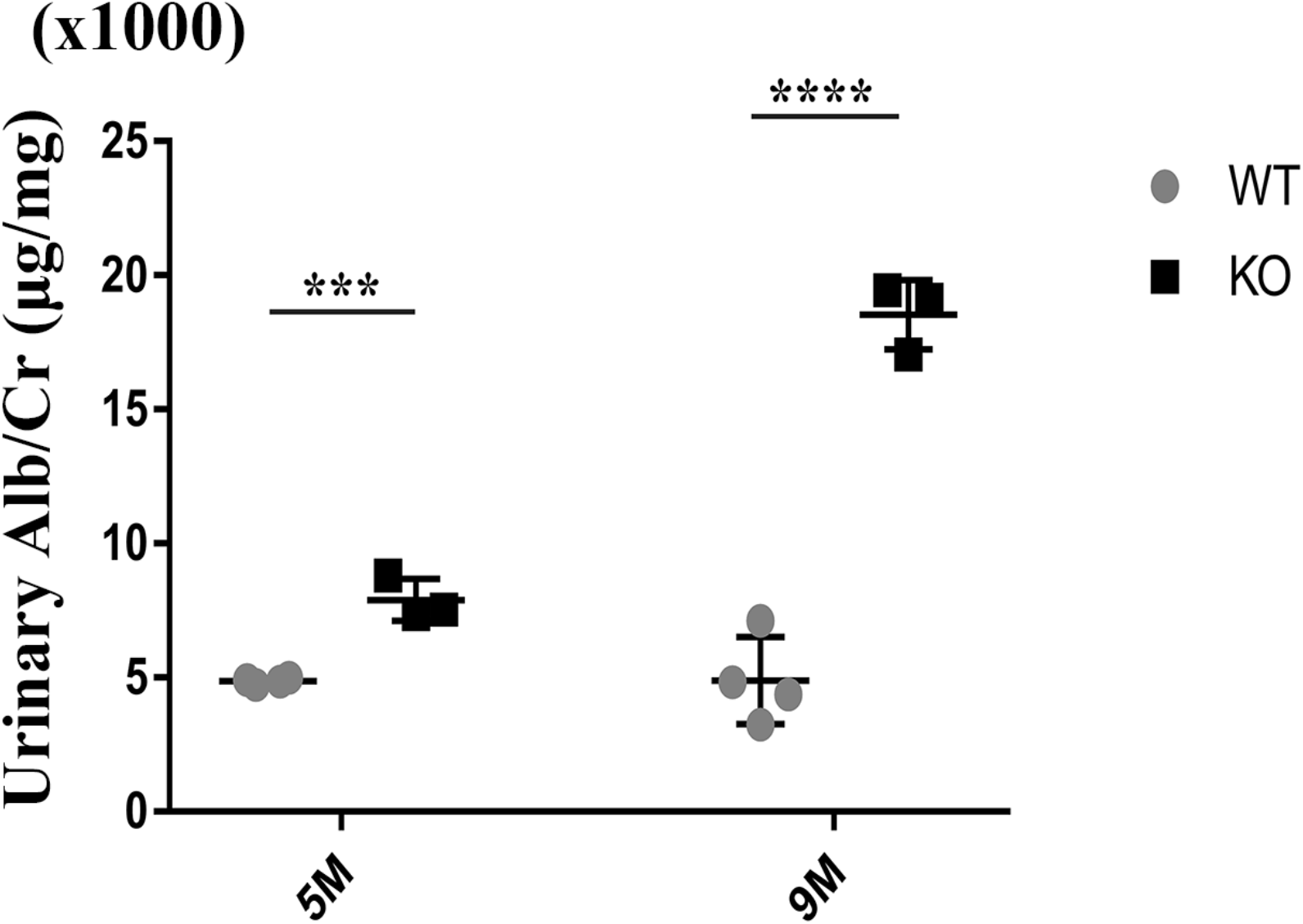
Deletion of *Tmem30a* in podocytes resulted in albuminuria. Urine biochemical analysis was performed in 5-month-old and nine-month-old WT and *Tmem30a* KO mice. Quantitation of urinary albumin in WT and *Tmem30a* KO mice showed that *Tmem30a* KO mice exhibited albuminuria at 5 months of age, which became severe by 9 months of age. Sample size, n=3 for both WT and KO mice. n= number of independent biological replicates. ***P<0.001, ****P<0.0001, ns=no significance. The data are represented as the mean ± SEM.

Albuminuria is an unambiguous symptom of the compromised integrity of the glomerular filtration barrier [24]. With increased protein passage from blood into urine, the proximal tubular reuptake mechanism is stimulated to reabsorb an increasing amount of protein until the reabsorption capacity is saturated [25]. Once the amount of protein excreted from blood exceeds the reabsorption capacity of the proximal tubule, albuminuria occurs. Mounting evidence indicates that albuminuria is one of the major features of various kidney diseases, or at least that albuminuria accelerates kidney disease progression to end-stage renal failure [26]. This indicates that defects in *Tmem30a* are a crucial cause of albuminuria.

### *Tmem30a* is essential for the survival and function of podocytes

*Tmem30a* deletion results in albuminuria, implying podocyte injury and loss in *Tmem30a* KO mice. We reasoned that this mouse model should allow us to address the question about the role of *Tmem30a* in the glomerular filtration barrier and progression of nephrotic syndrome. We next examined whether *Tmem30a* is required for the survival of podocytes. Paraffin sections from both *Tmem30a* KO mice and WT mice at 5 months and 9 months of age were subjected to immunostaining for WT1 and synaptopodin, which are two representative markers of differentiated podocytes. WT1 is the nuclear marker of differentiated podocytes used to assess the state of mature podocytes. In the kidney of *Tmem30a* KO mice, the number of WT1-positive cells in glomeruli was dramatically decreased by 5 months of age in a pattern consistent with the severity of diffuse glomerulosclerosis, indicating a loss of podocytes (Fig. 3a-b). Synaptopodin is an actin-associated protein that may play a role in actin-based cell shape and motility [27, 28]. Synaptopodin expression was also observed in the podocytes of WT mice but was hardly detectable in KO mice (Fig. 3c). The results of immunostaining for WT1 and synaptopodin confirm the loss of mature podocytes in *Tmem30a* KO mice, indicating that *Tmem30a* plays an essential role in the survival and function of podocytes.

**Fig. 3.**
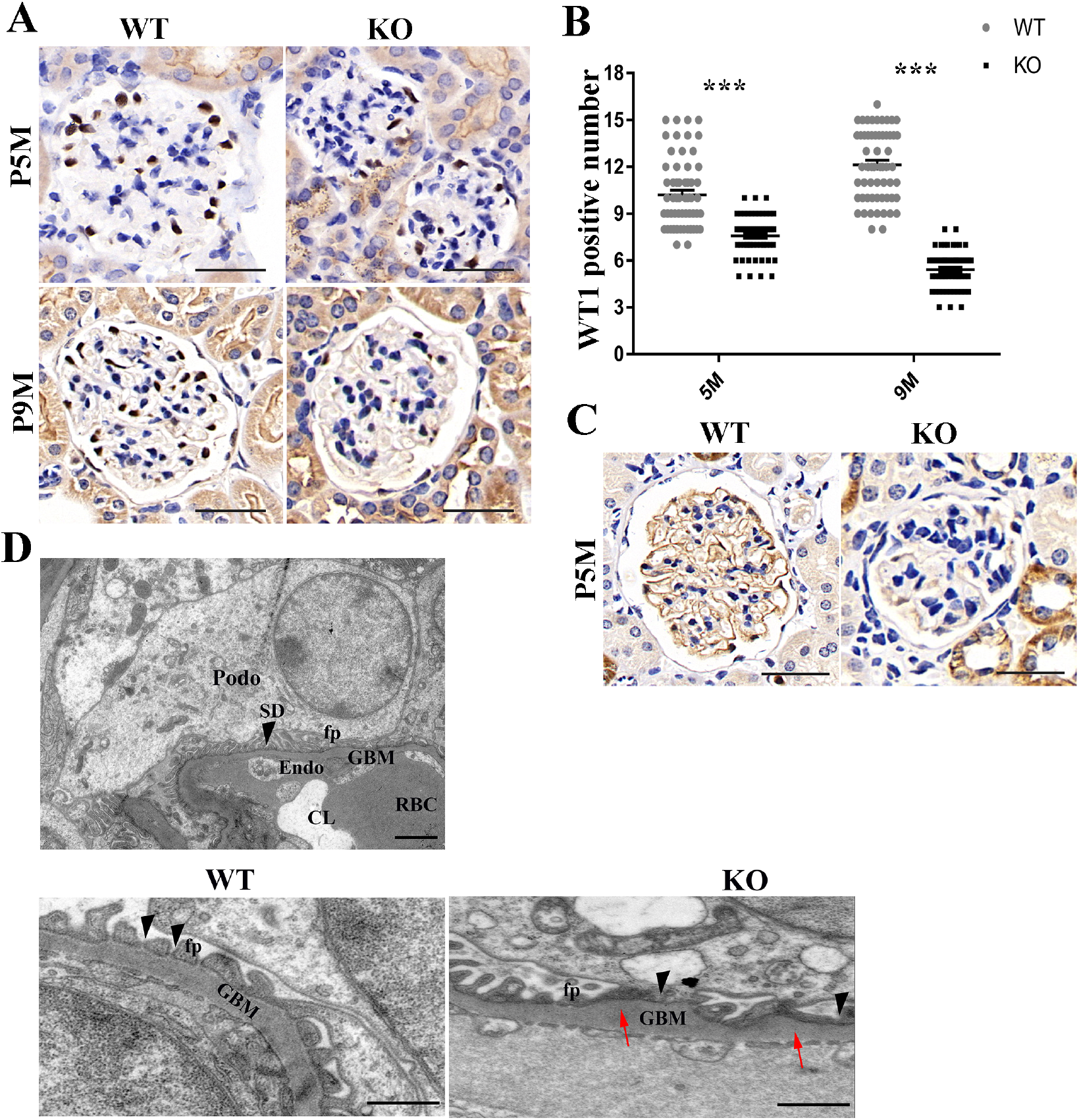
*Tmem30a* deficiency impaired podocyte survival and function. **(A)** Immunohistochemical staining of kidney sections revealed that the number of WT1-positive cells in glomeruli dramatically decreased after 5 months in the KO mice compared to the WT littermates, indicating podocyte degeneration in *Tmem30a* KO mice (scale bar: 50 µm). **(B)** Quantification of WT1-positive cells in the glomeruli of both WT and KO mice. n=200. Mean±SEM. ***P<0.001. **(C)** Immunohistochemical staining of paraffin-embedded kidney sections from *Tmem30a* WT and KO mice for synaptopodin revealed the loss of synaptopodin by 5 months of age. Positive staining for synaptopodin was hard to detect at 5 months, indicating podocyte loss. Scale bar: 50 µm. **(D)** Transmission electron microscopy images of glomeruli in *Tmem30a* WT and KO mice at 5 months. The upper panel shows normal glomerular filtration barrier and slit diaphragm (SD) formed between adjacent foot processes (fp) (scale bar: 2 µm). Compared with WT mice, KO mice exhibited increasing glomeruli base membrane (GBM) (red arrows), fusion of foot processes and lack of slit diaphragms (black arrowheads) (scale bar: 500 nm) .CL, capillary lumen; GBM, glomerular basement membrane; Endo, endothelium; RBC, red blood cell; Podo, podocyte; SD, slit diaphragm; fp, foot process.

To further examine the role of *Tmem30a* in FP formation, the ultrastructure in WT and KO mice at five months of age was analysed by TEM (Fig. 3d). *Tmem30a* WT mice showed normal podocyte, podocyte FP and GBM architecture (Fig. 3d, upper and lower left panel). In contrast, *Tmem30a* KO mice showed podocyte FP effacement, lack of a SD and increases in the GBM (Fig. 3d lower right panel), suggesting that *Tmem30a* deficiency causes impaired podocyte FP formation or imbalanced protein-protein interactions within the SD multiprotein complex, resulting in an impaired filtration barrier in the kidney.

### Loss of *Tmem30a* in podocytes causes endoplasmic reticulum (ER) stress

A previous study suggested that the loss of *Tmem30a* in Purkinje cells induced ER stress and subsequent progressive degeneration of Purkinje cells, demonstrating the vital function of *Tmem30a* in intracellular trafficking [15]. It is reasonable to suspect that podocyte injury and loss in *Tmem30a* KO mice is likely to induce ER stress. Western blot analysis showed that the expression of ER stress-related proteins, including CHOP and PDI, was upregulated in *Tmem30a* KO mice compared with WT mice at 5 months of age, indicating the presence of ER stress in KO podocytes (Fig. 4).

**Fig. 4.**
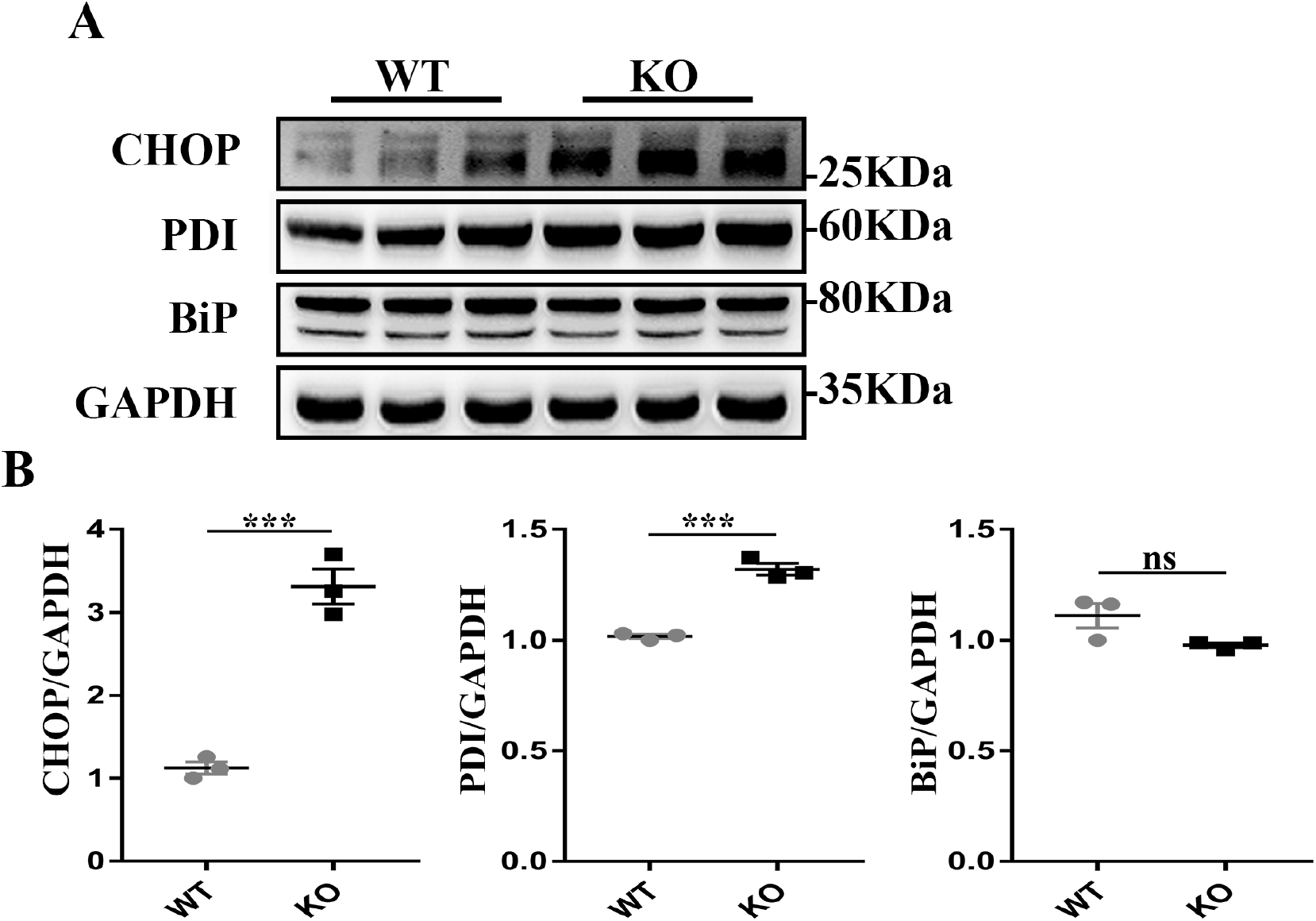
Loss of *Tmem30a* causes ER stress in podocytes. Western blot analysis of isolated glomeruli proteins in WT and *Tmem30a* KO mice at 5 months of age. (A) Western blotting was performed to detect the expression of CHOP, PDI and BiP, and GAPDH was probed as a loading control. (B-D) Quantitative analysis of blots. Sample size, n=3. ***, *P*<0.001. The data represent the mean±SEM.

### *Tmem30a* KO mice develop severe glomerulosclerosis

Kidney sections from both WT and KO mice at 2.5 months, 5 months and 9 months of age were analysed by light microscopy to assess pathological changes. PAS staining of kidney sections revealed normal nephrogenesis in *Tmem30a* KO mice, and the predominant renal changes were confirmed to be related to glomeruli (Fig. 5).

**Fig. 5.**
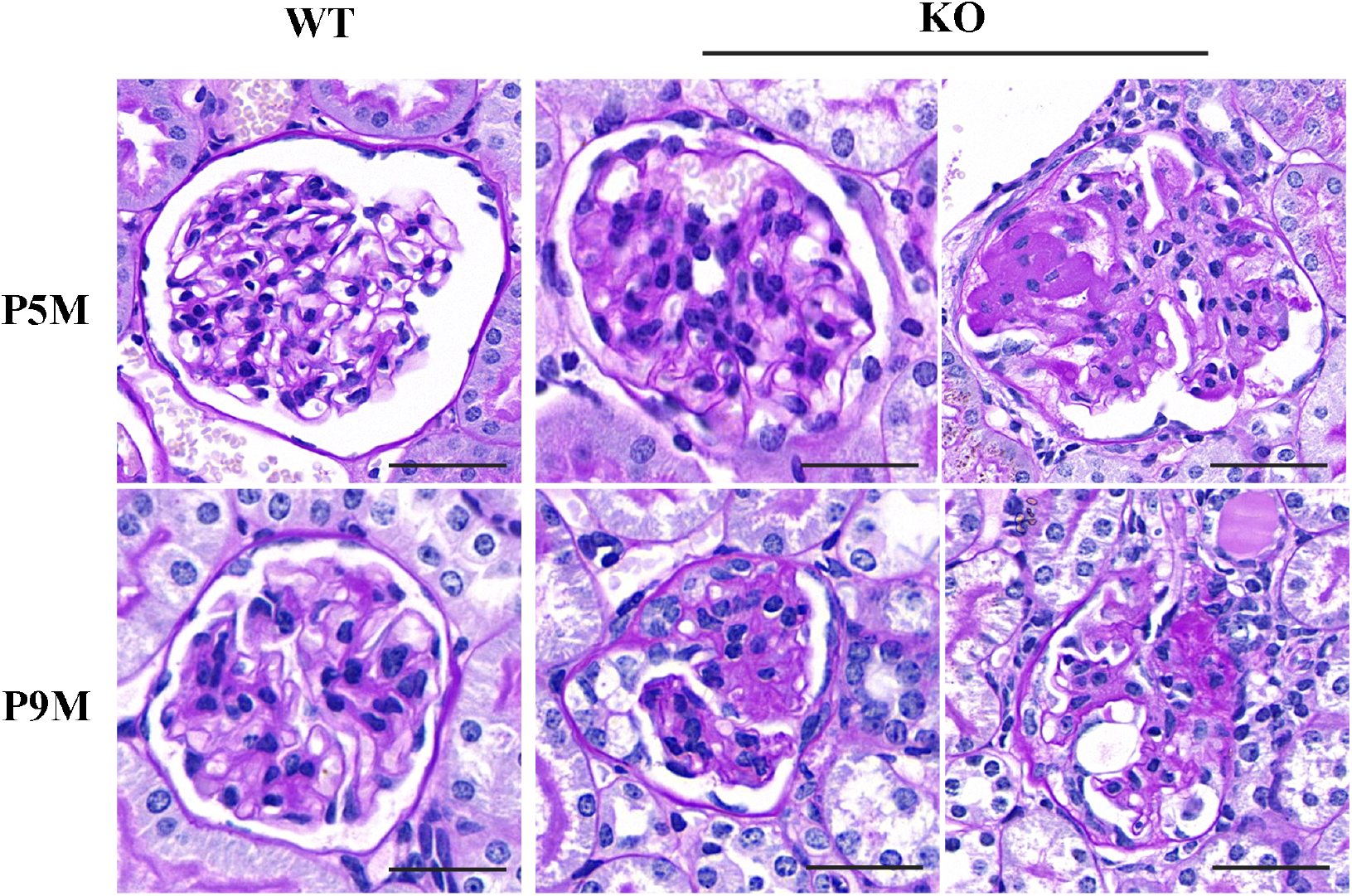
Glomerular sclerosis in *Tmem30a* KO mice. Representative light microscopy images of periodic acid-Schiff (PAS)-stained kidney samples from WT and KO mice. By the age of 5 months, glomeruli showed mesangial cell proliferation and increased extracellular matrix deposition with segmental glomerulosclerosis and (mild) adhesions to Bowman’s capsule in *Tmem30a* KO mice. At 9 months of age, more glomeruli were damaged and exhibited varying severities of pathological phenotypes as the disease progressed, such as mesangial cell proliferation and increased extracellular matrix deposition with segmental glomerulosclerosis (left panel of P9M KO) and adhesions to Bowman’s capsule (right panel of P9M KO). Sample size, n=3. Scale bar, 50 µm.

The size of the kidney in *Tmem30a* KO mice was generally the same as that of the kidney in WT mice (data not shown). Interesting, by 5 months, *Tmem30a* KO mice exhibited multiple pathologic processes, including slight and sever mesangial hyperplasia, mesangial cell proliferation with ECM deposition, capsular synechia and even glomerular sclerosis was visible throughout renal cortex (Fig. 5, upper panel). And by 9 months, more normal glomeruli were affected by loss of *Tmem30a* and showed prominent glomerular sclerosis (Fig. 5, lower panel). These data suggest that the kidney is undergoing a pathological process of FSGS, which also explains the absence of prenatal mortality.

### TMEM30A expression is reduced in patients with podocytopathy, including MCD and MN

TMEM30A is expressed in human glomeruli (Fig. 6 A). To evaluate the clinical importance of TMEM30A, we analysed the expression of TMEM30A in kidney samples from patients with podocytopathy (MCD, MN, and DN), samples from patients with IgAN and adjacent normal tissues from patients with renal tumours as controls (clinical information of the subjects in Table 1). Compared with the normal controls, the MCD and MN kidney sample showed significantly reduced TMEM30A expression levels (Fig. 6 B, C). Conversely, the expression level in tissue from IgAN patients showed no significant reduction. Although the expression of TMEM30A in tissue from DN patients showed no significant difference, it showed a downward trend. These data suggest that the expression of *TMEM30A* is decreased in podocytopathy, especially in MCD and MN, and that TMEM30A is essential for podocytes.

**Table 1.**
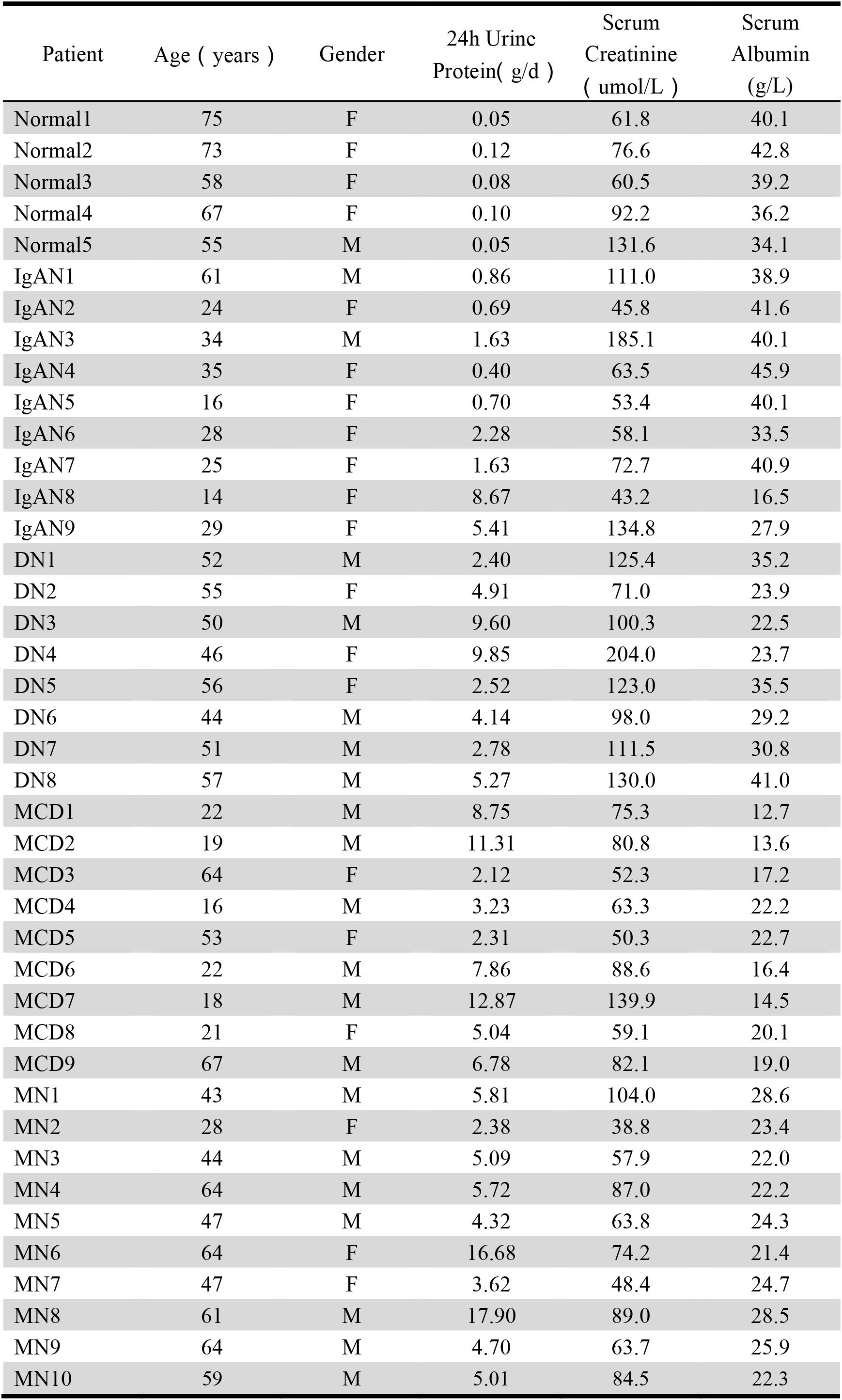
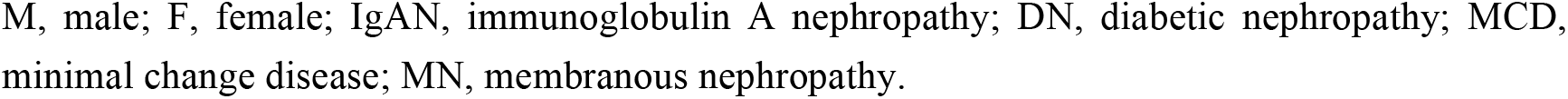
Baseline characteristics of the enrolled patients.

| Patient | Age ( years ) | Gender | 24h Urine<br>Protein( g/d ) | Serum<br>Creatinine<br>( umol/L ) | Serum<br>Albumin<br>(g/L) |
| --- | --- | --- | --- | --- | --- |
| Normal1 | 75 | F | 0.05 | 61.8 | 40.1 |
| Normal2 | 73 | F | 0.12 | 76.6 | 42.8 |
| Normal3 | 58 | F | 0.08 | 60.5 | 39.2 |
| Normal4 | 67 | F | 0.10 | 92.2 | 36.2 |
| Normal5 | 55 | M | 0.05 | 131.6 | 34.1 |
| IgAN1 | 61 | M | 0.86 | 111.0 | 38.9 |
| IgAN2 | 24 | F | 0.69 | 45.8 | 41.6 |
| IgAN3 | 34 | M | 1.63 | 185.1 | 40.1 |
| IgAN4 | 35 | F | 0.40 | 63.5 | 45.9 |
| IgAN5 | 16 | F | 0.70 | 53.4 | 40.1 |
| IgAN6 | 28 | F | 2.28 | 58.1 | 33.5 |
| IgAN7 | 25 | F | 1.63 | 72.7 | 40.9 |
| IgAN8 | 14 | F | 8.67 | 43.2 | 16.5 |
| IgAN9 | 29 | F | 5.41 | 134.8 | 27.9 |
| DN1 | 52 | M | 2.40 | 125.4 | 35.2 |
| DN2 | 55 | F | 4.91 | 71.0 | 23.9 |
| DN3 | 50 | M | 9.60 | 100.3 | 22.5 |
| DN4 | 46 | F | 9.85 | 204.0 | 23.7 |
| DN5 | 56 | F | 2.52 | 123.0 | 35.5 |
| DN6 | 44 | M | 4.14 | 98.0 | 29.2 |
| DN7 | 51 | M | 2.78 | 111.5 | 30.8 |
| DN8 | 57 | M | 5.27 | 130.0 | 41.0 |
| MCD1 | 22 | M | 8.75 | 75.3 | 12.7 |
| MCD2 | 19 | M | 11.31 | 80.8 | 13.6 |
| MCD3 | 64 | F | 2.12 | 52.3 | 17.2 |
| MCD4 | 16 | M | 3.23 | 63.3 | 22.2 |
| MCD5 | 53 | F | 2.31 | 50.3 | 22.7 |
| MCD6 | 22 | M | 7.86 | 88.6 | 16.4 |
| MCD7 | 18 | M | 12.87 | 139.9 | 14.5 |
| MCD8 | 21 | F | 5.04 | 59.1 | 20.1 |
| MCD9 | 67 | M | 6.78 | 82.1 | 19.0 |
| MN1 | 43 | M | 5.81 | 104.0 | 28.6 |
| MN2 | 28 | F | 2.38 | 38.8 | 23.4 |
| MN3 | 44 | M | 5.09 | 57.9 | 22.0 |
| MN4 | 64 | M | 5.72 | 87.0 | 22.2 |
| MN5 | 47 | M | 4.32 | 63.8 | 24.3 |
| MN6 | 64 | F | 16.68 | 74.2 | 21.4 |
| MN7 | 47 | F | 3.62 | 48.4 | 24.7 |
| MN8 | 61 | M | 17.90 | 89.0 | 28.5 |
| MN9 | 64 | M | 4.70 | 63.7 | 25.9 |
| MN10 | 59 | M | 5.01 | 84.5 | 22.3 |

- 1 M, male; F, female; IgAN, immunoglobulin A nephropathy; DN, diabetic nephropathy; MCD, - 2 minimal change disease; MN, membranous nephropathy. - 3 - 4

**Fig. 6.**
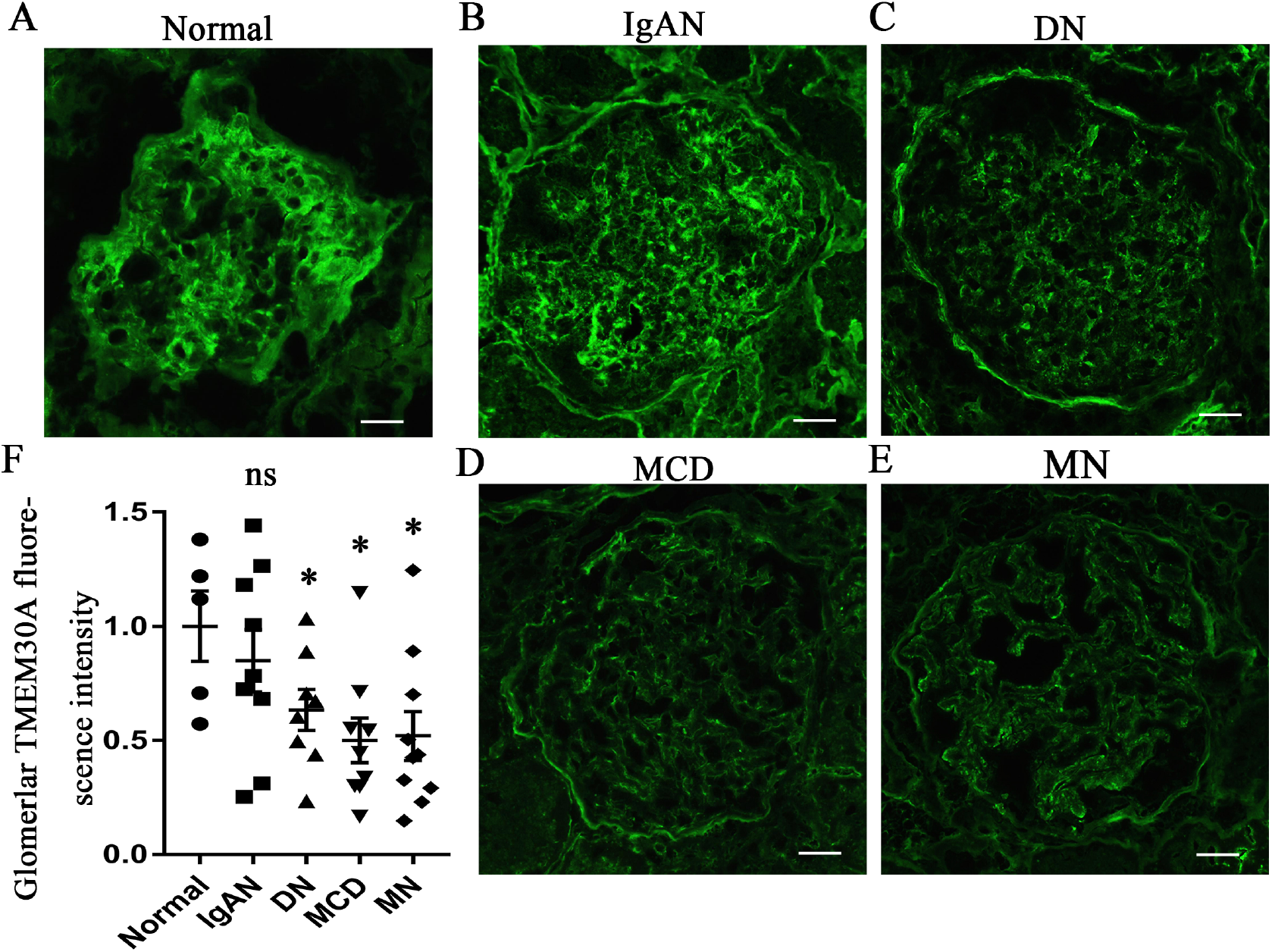
Immunofluorescence staining of human glomeruli revealed reduced expression of *TMEM30A* in minimal change disease and membranous nephropathy patients. (A) Immunofluorescence images of TMEM30A expression in normal human glomerular tissue. (B) Immunofluorescence images of TMEM30A expression in glomerular tissue from human patients with IgAN, DN, MCD and MN. (C). Quantification of the intensity of fluorescent staining for human glomerular TMEM30A. Mean±SEM. Normal group (n=5) vs IgAN group (n=9), P=0.455; normal group vs DN group (n=8), P=0.079; normal group vs MCD group (n=9), P=0.043; normal group vs MN group (n=10), P=0.019; * vs normal group, P<0.05. IgAN, immunoglobulin A nephropathy; DN, diabetic nephropathy; MCD, minimal change disease; MN, membranous nephropathy. Scale bar, 50 µm.

## Discussion

*The **β***-subunit of PS flippase *Tmem30a* is essential for generating and maintaining the asymmetrical distribution of phospholipids to ensure cellular signal transduction [8, 29-31]. In this study, we found that *Tmem30a* plays a vital role in maintaining glomerular filtration barrier integrity by generating a podocyte-specific *Tmem30a* KO mouse model. *Tmem30a* loss leads to podocyte injury and loss, albuminuria, mesangial cell proliferation with mesangial matrix accumulation and eventually glomerulosclerosis as the disease progresses.

Podocyte injury and loss are now recognized as initiating factors leading to glomerulosclerosis in the progression of multiple variants of kidney diseases, such as DN, IgAN and FSGS [32-36]. Podocytes are terminally differentiated cells that cannot repopulate after loss. Although a subpopulation of parietal epithelial cells can transform into podocytes, the capacity for regeneration appears to be limited and cannot compensate for the loss of podocytes [37-40]. Thus, podocyte injury and loss result in additional podocyte stress and ultimately glomerulosclerosis.

Given that *Tmem30a* plays a vital function in intercellular trafficking, we investigated the representative expression of the ER stress markers in isolated glomeruli: CHOP, PDI and BiP. The results showed upregulated expression of CHOP and PDI in *Tmem30a* KO mice, implying induced ER stress in *Tmem30a* KO mice due to the loss of *Tmem30a* in podocytes. We evaluated the hallmark of the impaired integrity of the glomerular filtration barrier, albuminuria, and found that *Tmem30a* KO mice showed albuminuria at 5 months after birth, indicating impaired podocytes. Albuminuria became more severe in *Tmem30a* KO mice at 9 months after birth (Fig. 2). The deletion of *Tmem30a* in podocytes resulted in a compromised glomerular filtration barrier at 5 months of age. The decreased immunostaining of synaptopodin was due to podocyte injury. In addition, TEM analysis further identified podocyte injury in *Tmem30a* KO mice: the intercellular junction and cytoskeletal structure of the FPs were altered, and the cells exhibited an effaced phenotype, indicating podocyte injury (Fig. 3d). SD structures disappeared, and albuminuria developed. Research on human kidney tissues showed decreased expression of *TMEM30A* in podocytopathy, especially in MCD and MN (Fig. 6), and validated the importance of TMEM30A in podocytes.

Mounting evidence suggests that mesangial cells are activated in numerous glomerular diseases and undergo proliferation and phenotypic alterations in response to glomerular injury, allowing glomerular structural recovery [41, 42]. However, compensatory activity after injury leads to the proliferation of mesangial cells along with abnormal ECM deposition, which results in glomerular fibrosis or sclerosis [43]. PAS staining of samples from *Tmem30a* KO mice at 5 months showed multiple pathologic process, approximately 12 out of ~200 glomeruli in *Tmem30a* KO mice exhibited mesangial cell proliferation, increased ECM deposition and even with segmental glomerulosclerosis. Furthermore, these pathological phenotypes became more severe and common at 9 months of age (Fig.4). These results indicate that glomerular disease caused by the lack of *Tmem30a* in podocytes progressed rapidly. It is possible that filtered macromolecules become trapped in the mesangium, causing the overreaction of mesangial cells and triggering an inflammatory response that plays a pivotal role in stimulating ECM synthesis, causing an imbalance between ECM synthesis and dissolution [44]. Persistent mesangial cell proliferation and ECM accumulation lead to glomerulosclerosis and end-stage rental failure.

In summary, our study reveals novel roles of *Tmem30a* in maintaining the integrity of the glomerular filtration barrier. The deletion of *Tmem30a* in podocytes resulted in podocyte degeneration, which led to a series of pathological phenotypic changes, including albuminuria, mesangial cell proliferation, mesangial matrix accumulation and glomerulosclerosis. One possibility is that *Tmem30a* deficiency causes defects in protein folding and transport in the ER, causing ER stress, which leads to podocyte injury and loss. Another possibility is that *Tmem30a* loss impairs lipid raft formation. The SD is actually a lipid raft with a multiprotein complex, in which dynamic protein-protein interactions maintain the SD as the final form of selective filtration. This provides us with another unique perspective to understand the mechanism of podocyte damage. Further investigation is necessary to elucidate the molecular signalling pathway in podocytes after the deletion of *Tmem30a*.

## Acknowledgements

This study was supported by grants from the Department of Science and Technology of Sichuan Province (www.scst.gov.cn, 2020JDYH0027 and 2018JZ0019, XZhu) and the Sichuan Provincial People’s Hospital scientific research fund for clinicians (30305031165P, LP). The funders played no role in the study design, data collection or analysis or manuscript preparation.

## Data availability statement

All data are included in the manuscript.

## References

1. Devaux, P.F., Static and dynamic lipid asymmetry in cell membranes. Biochemistry, 1991. 30(5): p. 1163–73.

2. Daleke, D.L. and J.V. Lyles, Identification and purification of aminophospholipid flippases. Biochim Biophys Acta, 2000. 1486(1): p. 108–27.

3. Dhar, M.S., et al., Mice heterozygous for Atp10c, a putative amphipath, represent a novel model of obesity and type 2 diabetes. J Nutr, 2004. 134(4): p. 799–805.

4. van der Mark, V.A., et al., The lipid flippase heterodimer ATP8B1-CDC50A is essential for surface expression of the apical sodium-dependent bile acid transporter (SLC10A2/ASBT) in intestinal Caco-2 cells. Biochim Biophys Acta, 2014. 1842(12 Pt A): p. 2378–86.

5. Liu, L., et al., Hepatic Tmem30a Deficiency Causes Intrahepatic Cholestasis by Impairing Expression and Localization of Bile Salt Transporters. Am J Pathol, 2017. 187(12): p. 2775–2787.

6. Mansergh, F., et al., Mutation of the calcium channel gene Cacna1f disrupts calcium signaling, synaptic transmission and cellular organization in mouse retina. Hum Mol Genet, 2005. 14(20): p. 3035–46.

7. van der Velden, L.M., et al., Heteromeric interactions required for abundance and subcellular localization of human CDC50 proteins and class 1 P4-ATPases. J Biol Chem, 2010. 285(51): p. 40088–96.

8. Folmer, D.E., et al., Cellular localization and biochemical analysis of mammalian CDC50A, a glycosylated beta-subunit for P4 ATPases. J Histochem Cytochem, 2012. 60(3): p. 205–18.

9. Bryde, S., et al., CDC50 proteins are critical components of the human class-1 P4-ATPase transport machinery. J Biol Chem, 2010. 285(52): p. 40562–72.

10. Takatsu, H., et al., ATP9B, a P4-ATPase (a putative aminophospholipid translocase), localizes to the trans-Golgi network in a CDC50 protein-independent manner. J Biol Chem, 2011. 286(44): p. 38159–67.

11. Takatsu, H., et al., Phospholipid flippase activities and substrate specificities of human type IV P-type ATPases localized to the plasma membrane. J Biol Chem, 2016. 291(41): p. 21421.

12. Coleman, J.A., M.C. Kwok, and R.S. Molday, Localization, purification, and functional reconstitution of the P4-ATPase Atp8a2, a phosphatidylserine flippase in photoreceptor disc membranes. J Biol Chem, 2009. 284(47): p. 32670–9.

13. Coleman, J.A., et al., Phospholipid flippase ATP8A2 is required for normal visual and auditory function and photoreceptor and spiral ganglion cell survival. J Cell Sci, 2014. 127(Pt 5): p. 1138–49.

14. Zhang, S., et al., TMEM30A deficiency in endothelial cells impairs cell proliferation and angiogenesis. J Cell Sci, 2019. 132(7).

15. Yang, Y., et al., Disruption of Tmem30a results in cerebellar ataxia and degeneration of Purkinje cells. Cell Death Dis, 2018. 9(9): p. 899.

16. Yang, Y., et al., Tmem30a deficiency leads to retinal rod bipolar cell degeneration. J Neurochem, 2019. 148(3): p. 400–412.

17. Zhang, L., et al., Loss of Tmem30a leads to photoreceptor degeneration. Sci Rep, 2017. 7(1): p. 9296.

18. Yang, F., et al., Deletion of a flippase subunit Tmem30a in hematopoietic cells impairs mouse fetal liver erythropoiesis. Haematologica, 2019. 104(10): p. 1984–1994.

19. Li, N., et al., Tmem30a Plays Critical Roles in Ensuring the Survival of Hematopoietic Cells and Leukemia Cells in Mice. Am J Pathol, 2018. 188(6): p. 1457–1468.

20. Tryggvason, K., J. Patrakka, and J. Wartiovaara, Hereditary proteinuria syndromes and mechanisms of proteinuria. N Engl J Med, 2006. 354(13): p. 1387–401.

21. Bose, B. and D. Cattran, Glomerular diseases: FSGS. Clin J Am Soc Nephrol, 2014. 9(3): p. 626–32.

22. Moeller, M.J., et al., Podocyte-specific expression of cre recombinase in transgenic mice. Genesis, 2003. 35(1): p. 39–42.

23. Stevens, M. and S. Oltean, Assessment of Kidney Function in Mouse Models of Glomerular Disease. J Vis Exp, 2018(136).

24. Coleman, J.A. and R.S. Molday, Critical role of the beta-subunit CDC50A in the stable expression, assembly, subcellular localization, and lipid transport activity of the P4-ATPase ATP8A2. J Biol Chem, 2011. 286(19): p. 17205–16.

25. Remuzzi, G., A. Schieppati, and P. Ruggenenti, Clinical practice. Nephropathy in patients with type 2 diabetes. N Engl J Med, 2002. 346(15): p. 1145–51.

26. Maunsbach, A.B., Albumin absorption by renal proximal tubule cells. Nature, 1966. 212(5061): p. 546–7.

27. Abbate, M., C. Zoja, and G. Remuzzi, How does proteinuria cause progressive renal damage? J Am Soc Nephrol, 2006. 17(11): p. 2974–84.

28. Asanuma, K., et al., Synaptopodin regulates the actin-bundling activity of alpha-actinin in an isoform-specific manner. J Clin Invest, 2005. 115(5): p. 1188–98.

29. Huber, T.B., et al., Bigenic mouse models of focal segmental glomerulosclerosis involving pairwise interaction of CD2AP, Fyn, and synaptopodin. J Clin Invest, 2006. 116(5): p. 1337–45.

30. Paulusma, C.C., et al., ATP8B1 requires an accessory protein for endoplasmic reticulum exit and plasma membrane lipid flippase activity. Hepatology, 2008. 47(1): p. 268–78.

31. Kato, U., et al., Role for phospholipid flippase complex of ATP8A1 and CDC50A proteins in cell migration. J Biol Chem, 2013. 288(7): p. 4922–34.

32. McConkey, M., et al., Cross-talk between protein kinases Czeta and B in cyclic AMP-mediated sodium taurocholate co-transporting polypeptide translocation in hepatocytes. J Biol Chem, 2004. 279(20): p. 20882–8.

33. Wiggins, R.C., The spectrum of podocytopathies: a unifying view of glomerular diseases. Kidney Int, 2007. 71(12): p. 1205–14.

34. Lemley, K.V., et al., Podocytopenia and disease severity in IgA nephropathy. Kidney Int, 2002. 61(4): p. 1475–85.

35. Petermann, A.T., et al., Viable podocytes detach in experimental diabetic nephropathy: potential mechanism underlying glomerulosclerosis. Nephron Exp Nephrol, 2004. 98(4): p. e114–23.

36. Kihara, I., et al., Podocyte detachment and epithelial cell reaction in focal segmental glomerulosclerosis with cellular variants. Kidney Int Suppl, 1997. 63: p. S171–6.

37. Vogelmann, S.U., et al., Urinary excretion of viable podocytes in health and renal disease. Am J Physiol Renal Physiol, 2003. 285(1): p. F40–8.

38. Lasagni, L., et al., Notch activation differentially regulates renal progenitors proliferation and differentiation toward the podocyte lineage in glomerular disorders. Stem Cells, 2010. 28(9): p. 1674–85.

39. Pippin, J.W., et al., Cells of renin lineage are progenitors of podocytes and parietal epithelial cells in experimental glomerular disease. Am J Pathol, 2013. 183(2): p. 542–57.

40. Zhang, J., et al., Podocyte repopulation by renal progenitor cells following glucocorticoids treatment in experimental FSGS. Am J Physiol Renal Physiol, 2013. 304(11): p. F1375–89.

41. Hackl, M.J., et al., Tracking the fate of glomerular epithelial cells in vivo using serial multiphoton imaging in new mouse models with fluorescent lineage tags. Nat Med, 2013. 19(12): p. 1661–6.

42. Johnson, R.J., et al., The activated mesangial cell: a glomerular “myofibroblast”? J Am Soc Nephrol, 1992. 2(10 Suppl): p. S190–7.

43. Samarakoon, R., et al., Induction of renal fibrotic genes by TGF-beta1 requires EGFR activation, p53 and reactive oxygen species. Cell Signal, 2013. 25(11): p. 2198–209.

44. Zhao, J.H., Mesangial Cells and Renal Fibrosis. Adv Exp Med Biol, 2019. 1165: p. 165–194.

45. Santini, E., et al., Effects of different LDL particles on inflammatory molecules in human mesangial cells. Diabetologia, 2008. 51(11): p. 2117–25.

